# Interpretable machine learning identifies paediatric Systemic Lupus Erythematosus subtypes based on gene expression data

**DOI:** 10.1101/2021.06.03.446884

**Authors:** Sara A. Yones, Alva Annett, Patricia Stoll, Klev Diamanti, Linda Holmfeldt, Carl Fredrik Barrenäs, Jennifer R. S. Meadows, Jan Komorowski

**Author notes:** Equal contribution (Jennifer R. S. Meadows, Jan Komorowski).

## Abstract

Transcriptomic analyses are commonly used to identify differentially expressed genes between patients and controls, or within individuals across disease courses. These methods, whilst effective, cannot encompass the combinatorial effects of genes driving disease. We applied rule-based machine learning (RBML) models and rule networks (RN) to an existing paediatric Systemic Lupus Erythematosus (SLE) blood expression dataset, with the goal of developing gene networks to separate low and high disease activity (DA1 and DA3). The resultant model had an 81% accuracy to distinguish between DA1 and DA3, with unsupervised hierarchical clustering revealing additional subgroups indicative of the immune axis involved or state of disease flare. These subgroups correlated with clinical variables, suggesting that the gene sets identified may further the understanding of gene networks that act in concert to drive disease progression. This included roles for genes i) induced by interferons (*IFI35* and *OTOF*), ii) key to SLE cell types (*KLRB1* encoding CD161), or iii) with roles in autophagy and NF-κB pathway responses (*CKAP4*). As demonstrated here, RBML approaches have the potential to reveal novel gene patterns from within a heterogeneous disease, facilitating patient clinical and therapeutic stratification.

## INTRODUCTION

Paediatric systemic lupus erythematosus (pSLE) is a rare, clinically and genetically heterogeneous systemic autoimmune disease with a prevalence of between 3.3-8.8 per 100,000 children^1^. The disease course is unpredictable, with periods of remission and flares that lead to cumulative damage over time^2^. SLE is classified by the presence of at least 4 out of 11 of clinical criteria^3^, with disease activity (DA) severity calculated based on composite scores, including Systemic Lupus Erythematosus Disease Activity Index (SLEDAI)^4^. Genetic studies have identified more than thirty genes associated with SLE, including those driven by interferons^5^, or those controlling inflammation and tissue response to injury^6^. Together these have been used to highlight the link between SLE and viral responses^7^. However, the trigger that initiates the expression of these genes and the progression of SLE disease remains poorly understood^8^.

Efforts to unravel the SLE gene expression pathway have been initiated. A 2016 study of paediatric disease examined the personal transcriptomic profiles of 158 patients using linear mixed models built on blood expression data from 15,386 transcripts^9^. The transcript panel utilised for this process considered each gene locus individually, and correlated the binary up- or down-regulation patterns with patient phenotypes. The result was the stratification of patients into distinct subclasses, with an enrichment of neutrophil expressed transcripts noted as a patient passed from the low DA1 state to the high DA3 form of disease. While the molecular pathways proposed by the study have led to a better understanding of personal disease progression^9^, the analysis lacked the co-predictive power of rule-based machine learning (RBML) models.

Machine learning (ML) approaches are well suited to address this process, as they can model and characterise data with very high dimensionality, such as that generated through personal transcriptomics.

However, the majority of methods work as black boxes. These offer little to no explanation in terms of how, and why, a specific classification decision is made. For clinical -omics, understanding how a classification decision is made, may offer insight into the underlying biological mechanisms, for example contrasting a disease state to healthy controls^11^. Interpretable ML methods such as RBML models, offer classification transparency^11,12^. We applied RBML that is based on rough set theory. It uses Boolean reasoning to identify the minimal set of features that can discern decision classes (reducts). Reducts are subsequently overlaid onto transcriptomics data samples to create IF-THEN rules. One of the main advantages of this method is co-prediction, i.e., the identification of descriptors that collaboratively correctly classify samples from the data. Co-prediction can provide insight into the candidate biological processes beyond of what can be learnt through co-expression networks.

In the current study, we apply a RBML approach using rough sets to existing pSLE blood transcriptome data^9^. Here, the goal was to identify the genes and interactions that demarcate a low pSLE DA1 state from a high DA3 state. The disease sub-groups discovered were intersected with available clinical data, revealing gene sets key to the progression of disease and the involvement of the innate and acquired immune arms. These genes, and their protein products, have the potential to be translated to biomarkers, or could be suggested points for therapeutic intervention.

## RESULTS

### Minimum gene set model discerns DA1 from DA3

The initial rule-based model was built with R.ROSETTA^13^ using data from 629 unique patient clinical visits (observations) and the discretised gene expression value for each DA1 and DA3 patient visit (features: 33,006 probes for 629 observations; Figure 1). This initial model had an overall prediction accuracy of 71% using 10-fold cross validation (Supplementary Fig S1 online). The observations (visits) incorrectly classified by the model (Supplementary Fig S2 online) were pruned to achieve a better separation between DA1 and DA3 then intersected with the patient metadata in order to understand the potential reasons behind their misclassification. Observations were more likely to be pruned or removed based on patient treatment, low SLEDAI score or the number of days since diagnosis (Logistic regression p-value for all <0.05; Supplementary Fig S3 online). No significant association was observed between removed observations and clinical symptoms, and the significant associations were a reflection of observations removed from class DA3 (38%, 125/330 removed) rather than reductions from DA1 (53%, 157/299 removed).

**Figure 1.**
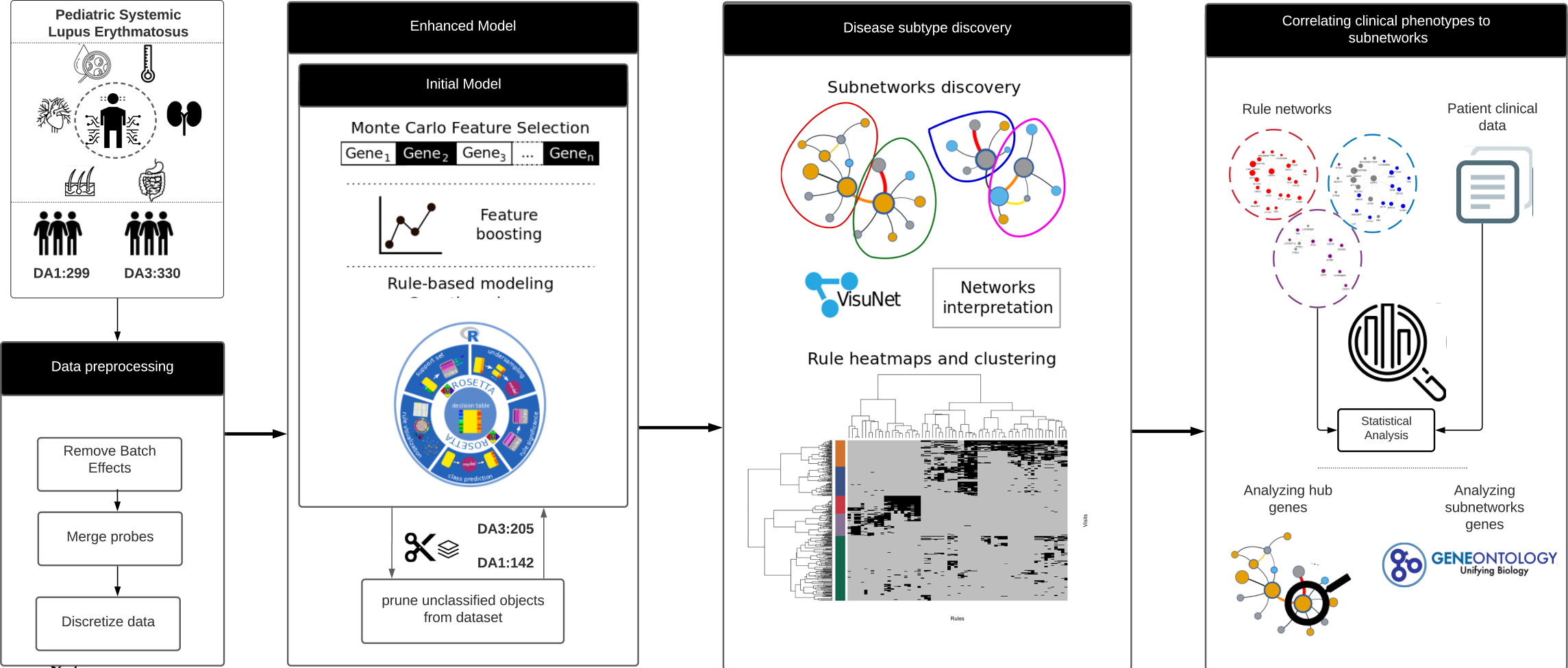
Overview of the modelling process implemented to classify and interrogate gene expression relationships between DA1 and DA3.

Following Monte Carlo Feature selection (MCFS)^14^ on the pruned dataset, 4,980 genes were available and subsequently used to build an enhanced rule-based model. Gene set enrichment analysis revealed terms connected to neutrophils (e.g., activation, mediation, degranulation) and the production and degradation of gene products (e.g., transcription initiation and nonsense-mediated decay; Supplementary Fig S4 online). This suggests a difference in neutrophil mediated immune response between patients with DA1 and DA3, a known functional shift in SLE manifestation between disease states^4^.

Feature boosting was performed to identify the optimal number of genes for the model (Figure 1). Empirical studies revealed that model accuracy was lost if more than 200 of the top 4,980 MCFS ranked genes were used for this process (Supplementary Fig S5 online). Iterative R. ROSETTA computational rounds added genes from the starting set of 200, with maximum model accuracy of 81% achieved with a minimum set of 34 genes (Figure 2; Supplementary Table S1 online). These genes were used in 22 and 44 classifying rules for DA1 and DA3 respectively. The model mirrored the structure of the initial model (Supplementary Fig S1 online). Figure 2 shows DA1 and DA3 were again split, however with a reduction of complexity, in terms of rules (edges) connecting the genes (nodes) and a refinement of the central hub genes. The 10% gain in the model accuracy provided improvement in terms of a clearer and visible separation between the disease activities in the rule networks (RN); this gain in accuracy was too small to imply an overfitting of the model. The similarity between the network of the initial model and the enhanced model implied that removed objects were unnecessary for classification of DA1 and DA3 since their removal did not significantly impact the main network structure or the rule model.

**Figure 2.**
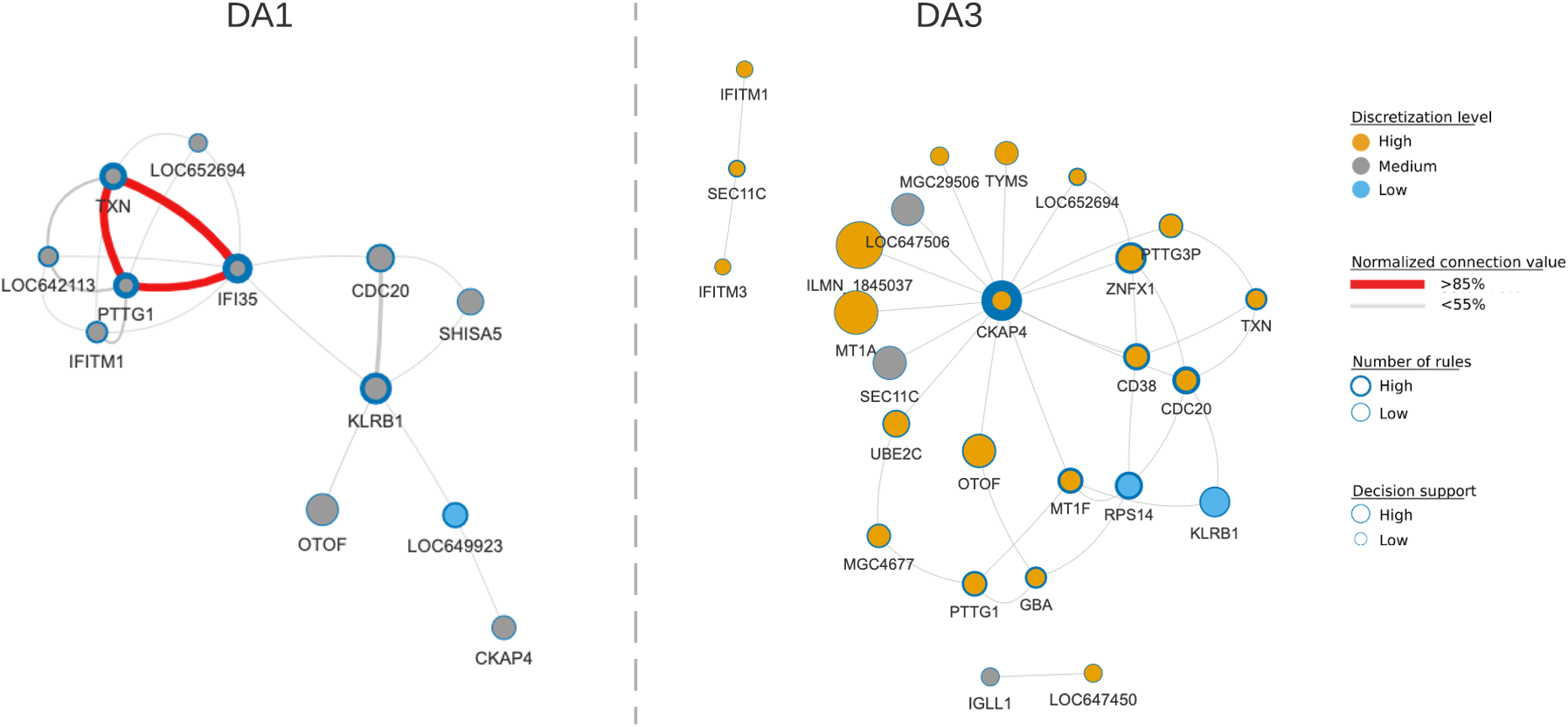
The rule networks discern the disease states. DA1 is largely defined by medium gene expression, whereas DA3 includes more genes, and those that were highly expressed. For each decision class, internal node colour indicates discretised gene expression value (high, medium, low; orange, grey, blue), node size is proportional to the number of objects supporting rules associated to a node, node border thickness is proportional to the number of rules associated to a node (low, high; circle border thin, thick) and edges connecting nodes represent normalised connection values (<55%, ≥85%; grey, red with increasing line thickness per support interval). The latter is the strength of the co-appearance of connected nodes in rules supporting a decision class. The network was filtered to visualise rules with minimum support of 10% and rule p-value ≤ 0.05.

In DA3, hub gene *CKAP4* was surrounded by a thick blue border, indicating the importance of this gene to predicting this disease state. In fact, *CKAP4* was a member of 14/44 co-prediction rules (Supplementary Table S1 online). The protein product of this gene, CKAP4 formerly CLIMP63, can act to regulate endoplasmic reticulum (ER) nanodomain homeostasis via shaping the luminal space or through interaction with other ER-resident proteins^15^. *CKAP4* was highly expressed (orange), whereas connected gene *SEC11C* showed a medium level of discretised expression (grey), and *RPS14* was lowly expressed (blue). In DA1, *IFI35* and *KLRB1* were both hub genes with medium expression levels. However, the latter had larger number of observations supporting its membership to rules (larger node size) but contributed to slightly fewer rules than *IFI35* (thinner circle border size: *IFI35*, 6/22 rules; *KLRB1*, 4/22 rules). CD161/NKR-P1A, encoded by *KLRB1*, is a surface receptor of natural killer (NK) cells and subtypes of T lymphocytes, whereas *IFI35* encodes the Interferon-induced 35 kDa protein, a proinflammatory damage-associated molecular pattern (DAMP) molecule in the innate immune pathway^16^.

The membership of genes to the rule networks was not discrete. For example, both *IFITM1*, which encodes interferon-induced transmembrane protein 1, and *KLRB1* appeared in DA1 and DA3 although with different expression values (Supplementary Table S1 online). The sharing of genes across rules was more common in DA1 (4/12 plotted genes are unique to DA1) than DA3, where 18/26 were unique to that class. The model showed that the type 1 interferon response term was limited to DA1 (*IFI35* and *IFITM1*) whereas B-cell activation was restricted to DA3 (*CD38* and *IGLL1*) (Supplementary Fig S6 online). However, while each term was enriched based on very few genes, it should be noted that these genes were present in multiple rules.

### Patient subgroups reflect clinical manifestations

Hierarchical clustering of the enhanced model results (i.e., the membership of observations for each rule) revealed five subgroups largely contained within DA1 (C1 and C2) or DA3 (C3, C4 and C5) (Figure 3A). The model and sub-groups were tested for significance to confirm that they cannot be attained by using random data. The significance was tested using permutation of DA state (p-value ≤ 0.05). These sub-clusters were subsequently projected onto the RNs (Figure 3B). Of note, the C1 and C2 sub-clusters were not restricted to the DA1 rule set, however C4 and C5 reflected partially intersecting networks that were all included in C3 and limited to DA3 (Figure 3B). In comparison to C3, the DA3 hub gene *CKAP4* was absent from C4, whereas the two small unconnected DA3 networks were absent from C5. Due to the small number of genes available for consideration, a sub-cluster-based gene enrichment analysis was not informative for all sets. The C4 and C5 enrichments were largely based on the combination of two genes (*MT1F*, *MT1A*; 11 genes available) and suggested response to ion levels (see Supplementary Fig S7 online), whereas C3 was again led by a small number of gene combinations (e.g., cell cycling and division: *CDC20*, *PTTG1*, *PTTG3P*, *UBE2C*; B-cell pathways: *CD38*, *GBA*, *TYM*) but this cluster also included the *MTIF*, and *MT1A* signals.

**Figure 3.**
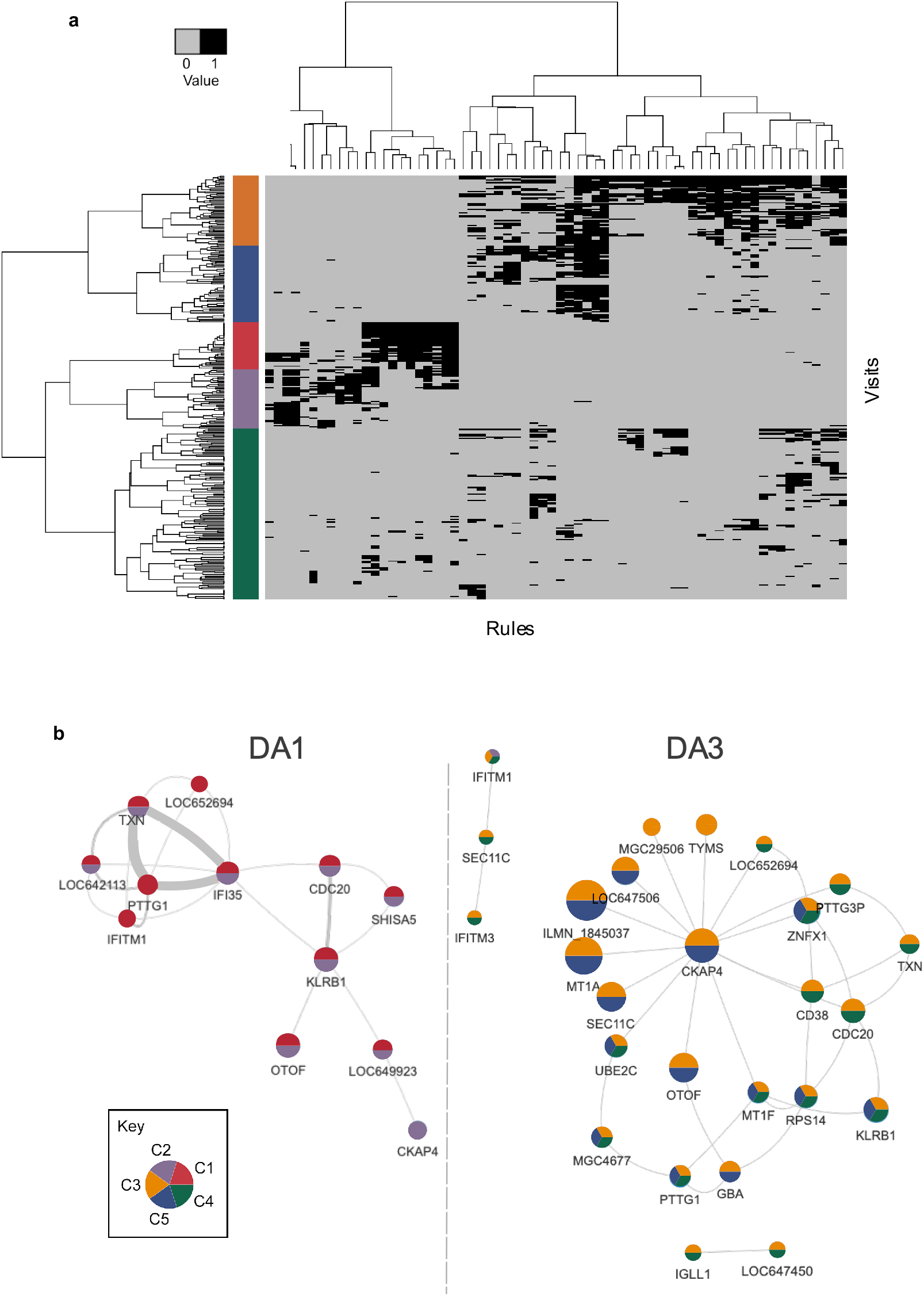
Hierarchical clustering of the model rules showed the major subdivision between the DA clusters. **(a)** Supported rules (black) and unsupported rules (grey) distinguish five disease subgroups that were projected into the **(b)** RN where group (cluster) membership is indicated by pie colour.

The relationship between clinical phenotype (Supplementary Table S3) and sub-cluster was explored in two ways. First, to assess clinical association to a sub-cluster, the phenotype values supporting that sub-cluster were compared to those that did not. To interrogate which rule(s) were driving that pattern, a similar assessment was performed, this time for visits supporting a rule within the sub-cluster. The examination of continuous phenotypes showed that these measures were only significantly different between the three DA3 clusters and not between the two DA1 clusters (Tukey HSD adjusted p-value <0.05; Supplementary Table S2 online). However, for DA1, the C1 and C2 clusters did contain the majority of low SLEDAI score visits (~1.7 in each, Supplementary Table S2, Supplementary Fig S8 online), with C1 tending towards lower alanine aminotransferase (ALT) and serum creatinine (CR) values compared with the C2 cluster. As expected, the DA3 cluster contained the higher SLEDAI scores (C4 ~8.8, C5 ~12.1, C3 ~14.6). C4 was largely reflective of low measures for anti-dsDNA antibody, erythrocyte sedimentation (ESR) and white blood cell count (WBC). C5 presented lower ALT and aspartate aminotransferase levels (AST), while C3 was most representative of active disease, with low complement factor C3 and C4 values (Supplementary Table S2, Supplementary Fig S8 online). Only two phenotypes, lymphocyte percent (LP) and neutrophil percent (NP), were significantly different in all pairwise DA3 cluster comparisons. LP was highest in C4 and NP, highest in C5. C3 was intermediate for both (Supplementary Table S2, Supplementary Fig S8 online).

In terms of categorical phenotypes, no significant association was detected between sex or race for each of the five clusters. In C1, the alopecia category was enriched when compared with all others (Fisher exact test p-value = 0.04; Supplementary Fig S8 online). In C2, the musculoskeletal term and both oral steroid and nephritis treatment groups were enriched (all Fisher exact test p-value < 0.05). Treatment could not be ruled out as the factor driving differences between this and other clusters (Supplementary Fig S9 online).

### Rules reveal which gene co-predictions drive phenotype correlation

To interrogate which genes and rules drove the phenotypic associations, a closer examination of the rules within the clusters was performed. To associate rules to the discovered clusters a frequency distribution was built for all rules with support set matching at least 10% of the visits assigned to each of the discovered clusters. Based on the distribution 20% match was an empirical threshold for assigning rules to each (Supplementary Fig 10 online). Figure 4 illustrates the fraction of rules from each cluster that were significantly associated with a phenotype, either continuous or categorical. Overall, rules from C1 or C3 were significantly associated with all phenotypes displayed (Figure 4), an enrichment not seen with the other clusters. Interestingly, whilst no individual continuous phenotype was significantly different between the two DA1 clusters, or categorical phenotype different between the DA3 clusters, the graphs clearly showed that the same was not true for the proportion of rules significantly associated with a phenotype in either class.

**Figure 4.**
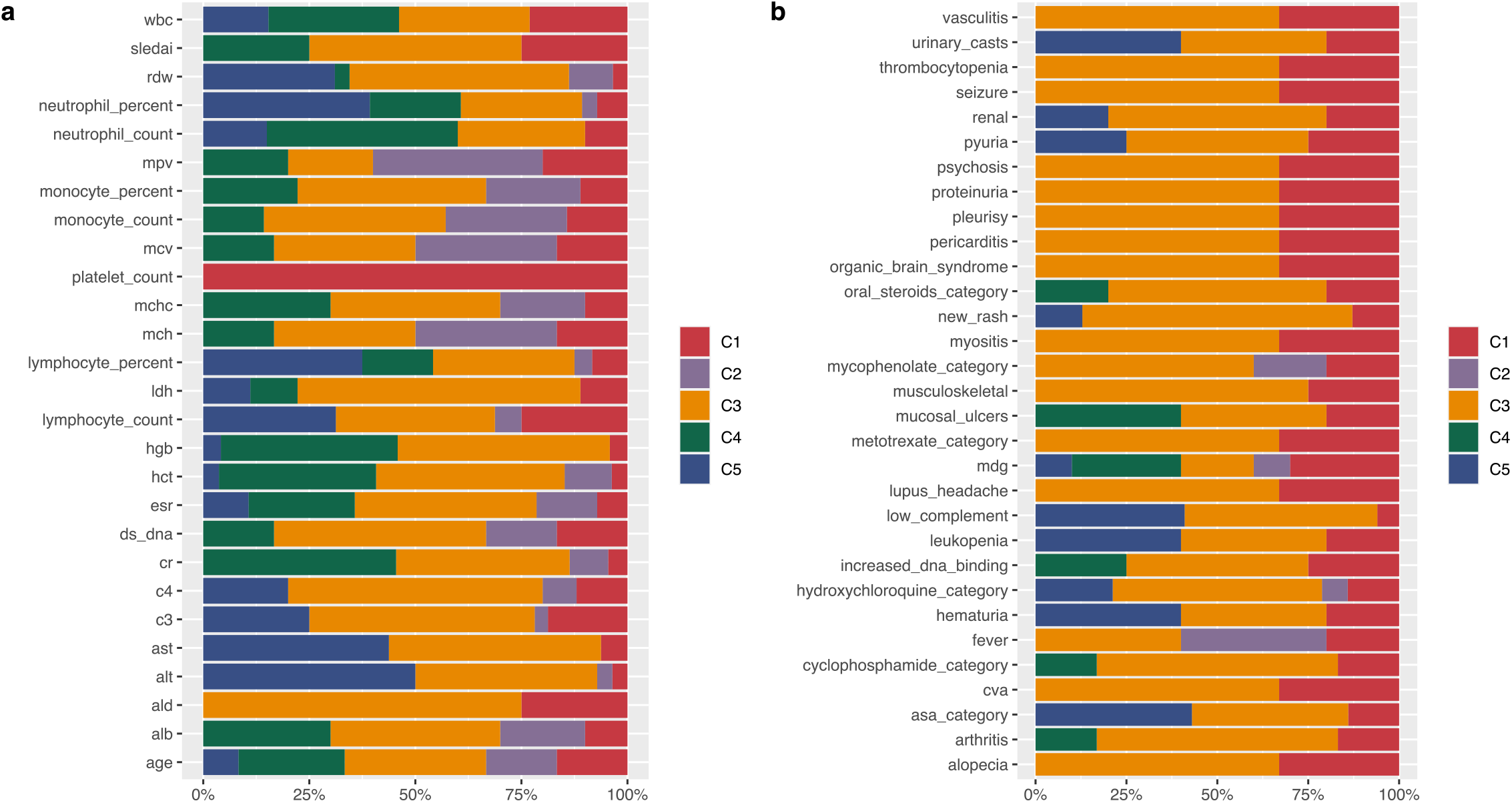
Fraction of rules per cluster significantly associated with **(a)** continuous and **(b)** categorical phenotypes. See Supplementary Table S3 online, for a list of clinical variables and phenotypes abbreviations.

For example, in the continuous class, rules from both DA1 clusters were significantly associated with lymphocyte count (LC; Figure 4A). There, three unique rules were contributed by C1 (rules 5, 44, 56), whereas the fourth rule was shared by both clusters (rule 41: *KLRB1*, *SEC11C*; Supplementary Table S4 online). Interestingly from the seven genes contained across the four rules, only the gene encoding the signal peptidase complex catalytic subunit, *SEC11C*, showed decreased expression, all others had medium values. This maintenance of gene expression likely explained the overall lack of significant difference between clusters for this trait.

For the DA3 clusters, a significant difference was recorded for the complement factor C3 phenotype between the C3 cluster (mean 62.1 mg/dL) and the C5 cluster (mean 85.9 mg/dL) (Wilcoxon test p-value = 2.4×10^−3^; Supplementary Table S2 online). An examination of the rules associated with phenotype C3 revealed that 17 rules were significantly linked to this phenotype in cluster C3, whilst only eight were found in the C5 cluster (Supplementary Table S4 online). All C5 rules were shared with C3, and no rules were contributed from C4 (Figure 4A). As expected from the associated RN, none of the nine rules unique to cluster C3 showed discrete gene membership, rather they served to illustrate how in comparison to C5, rules represented by network edges could introduce additional unique features that may serve to explain the phenotypic difference. For example, shared rule 15 (*CKAP4*, *MT1F*) can form an extended connection with rules 4 (*MT1F*, *KLRB1*), 23 (*CKAP4*, *SEC11C*) and 51 (*MT1F*, *PTTG1*), widening this network to include genes *KLRB1*, *SEC11C* and *PTTG1* (Supplementary Table S5 online). Each of these genes had previously been associated with SLE, but the link was not always clear. As noted before, *KLRB1*, expressed by NK cells and shown to be in the medium discretised expression level here, has been implicated in the regulation of the interferon gamma immune response^17^. *SEC11C*, encodes a subunit of microsomal signal peptidase complex and was the only DA3 gene maintained within medium levels for this phenotype. This gene was previously shown to be significantly down regulated in the T cells of adult SLE patients with low complement levels^17^. *PTTG1* was previously linked to SLE via SNP association^18^, although it was later shown that the risk allele was tagging the nearby microRNA, miR-146a, and this was down-regulated in European disease^19^.

## DISCUSSION

The use of machine learning in the current study has served to identify the key regulatory networks that underlie two disease states, DA1 and DA3, of the highly heterogeneous condition, paediatric systemic lupus erythematosus (pSLE). In doing so, the high dimensionality of data drawn from 33,006 gene expression measures across 629 paediatric patient visits has been reduced to co-predictive networks linked via genes. These genes were under-represented or down-weighted in published studies of SLE differential gene expression (DGE) profiling (Supplementary Fig S11 online). The result here was five sub-networks; two distinguishing DA1, perhaps as a result of treatment response, and three subgroups not related to treatment, within the more severe DA3 disease state.

Two major factors underpinned the difference in the results observed here, versus those generated by others in the field. The first was the study of patient visits, rather than individuals over time via longitudinal gene expression. The second was methodological, as RNs are co-predictive and as such, are conceptually different from co-expression networks. The goal here was to delve into the co-predictive RNs based on gene expression at different stages of disease, potentially creating a set of biomarkers, which could be used to stratify patient subgroups for clinical trials or personalised medicine based on their disease state at a particular time. This contrasts to the prognostic goals of others using the same dataset^9,20^.

Let us set the scene. For the transcriptomic data analysed here, the nodes of an RN are genes and their discretised expression values. The edges between two nodes of an RN are formed from pairs of genes and their discretized expression values as they co-occurred in the IF-part of rules (Figure 2). Significantly, in one outcome a gene may have one discretised value, but in the other outcome it will have a different value. It follows that each outcome has its own network. As such, co-prediction can provide insight into the candidate biological processes characteristic of the given outcome. For example, one combination of descriptors, i.e., pairs of gene and their discretised value may be associated to DA1 state, and another pair to DA3. This is in contrast to co-expression networks that identify genes that are co-expressed, not necessarily co-predictive of the outcomes.

SLE is a condition that spans the axes of both autoinflammatory and autoimmune disease. In this study, three DA3 subgroups were identified. The C3 sub-group sits on the autoimmune side, and had the clinical hallmarks of hypocomplementemia (low C3 and C4 clinical measures) in combination with high anti-dsDNA values, whilst the C4 sub-group likely represented the autoinflammatory side, with normal complement levels and low anti-dsDNA values (Supplementary Table S2 online). This was reinforced by the higher SLEDAI scores observed in C3 versus C4. Cluster C5 likely represented the intermediate stage between C3 and C4, where a significant shift between neutrophil and lymphocyte involvement is observed. This could translate to an immune complex driven disease state in C5, where the type I interferon process was active (low lymphocyte percent and increased neutrophil involvement). In studies using independent patient groups, both changes in complement ratio (C3/C4)^21^ and the categorisation of neutrophil to lymphocyte ratio (NLR)^22^, have been suggested as ways to distinguish SLE patient groups. Here, network analysis and unsupervised clustering combined both C3/C4 and NLR biomarker sets and resulted in three separate groups spanning these factors. The novelty in the current study lies in linking the clusters to co-predictive RNs, and this was the second major factor differentiating this work from others.

While the application of machine learning approaches to the big data sets generated by biology -omics is not new^23^, the approach used here removes the ‘black box’ interpretation of both the modelling and the results. This is required in the trade-off between predictability and interpretability^24^. Here we accepted the potentially reduced, but still high prediction accuracy of 81%, in favour of transparent classical models that perform well when the number of features available in the dataset (i.e. observations versus genes) outnumber the observations^25^. It is important to note that the rough sets approach to constructing rules is based on finding the minimal subsets of features that preserve discernibility of the decision classes from the original set. The rules will contain conjunctions of genes that may reflect different levels of gene regulation but that do not need to be co-expressed. In RNs, the genes and their regulation levels are associated to the outcome and discern the decision classes (here DA1 or DA3) based on the training data, while in co-expression networks the genes are co-expressed with other genes and may not discern the outcomes. The R.ROSETTA method used for constructing the model has been shown to outperform other existing rule based methods^25^, and has the key distinction of being the only method that can compute a significance level for the rules in the model. This is useful for calculating model prediction reliability, but it is the use of a minimum set of significant rules that served to highlight the genes contributing most strongly to the separate networks.

In practice, this was illustrated by the hub genes for DA1 (e.g., *IFI35*, *KLRB1*) and DA3 (e.g., *CKAP4*, *OTOF*; Figure 2). *IFI35* expression is stimulated in response to IFN-α/γ^26^ and it can act intracellularly as a negative switch in the innate immune pathway via retinoic acid-inducible gene I regulation^27^.

Extracellularly, the opposite effect has been observed, and the IFI35 molecule can act as a DAMP, and serve to activate the NF-κB pathway in macrophages via TLR4 signalling^16^. The end result is the release of proinflammatory cytokines, including interleukin 6 and tumour necrosis factor^16^. In DA1, *IFI35* expression is observed within the medium range, but a change in this value could be key in driving DA1 patients back to a remissive or inactive SLE state. Likewise, the maintained medium expression of *KLRB1* (encoding the surface receptor CD161) suggests a role for other cell sets, including natural killer (NK) cells and T lymphocytes in this lower disease state. The cell population expressing CD161 has been shown to be lower in SLE patients versus controls^28^. This is intriguing as this receptor can mark the NK cells that respond to innate cytokines and so promote innate inflammation^29^. Here again we see a contradiction between the promotion and reduction of the innate immune response.

While *CKAP4* was shown as a highly expressed hub gene in DA3, the protein product is most often reported to have a role in cancer, for example acting with RBP1 to induce autophagy in murine models of oral squamous cell carcinoma^30^. Autophagy can also play into the pathogenesis of SLE in a number of ways. Dysregulated autophagy can affect the regulation of T and B cell populations^31^, and increased autophagy can promote the NF-κB pathway response^32^. Through its interaction with ER-resident proteins, CKAP4 also has the potential to regulate or reflect the current state of cellular immune signalling^15^. For the individuals studied here, increased levels of CKAP4 may not be driving disease, but the finding opens a potential line of anti-CKAP4 antibody drug development for SLE patients; an avenue previously only promoted for cancer treatment^33^. Another DA3 hub gene, *OTOF*, is an interferon inducible gene, and has been recognised by others as a marker for SLE disease flares^34^. This is in keeping with the finding of *OTOF* in the C3 and C5 clusters, but not in C4. Recently it was suggested that through interaction with melatonin, OTOF may have a role in proteasome inhibition^35^, and so could function in the downstream signal transduction pathway of NF-κB^36^. While that study was focused on neuronal survival driven by melatonin ubiquitin proteasome system inhibition, a protective anti-inflammatory role of melatonin in SLE pathogenesis has been reported previously^37,38^. Gene networks acting through the fulcrum of *OTOF* may help to explain this action, and suggests that further investigation of melatonin treatment in SLE flare could be warranted.

The current analysis aimed to explore the different networks that underlie pSLE disease states with the goal of developing a minimum set of rules that could discern disease states DA1 from DA3. It is worth to mention that we did not aim to model the entire spectrum of pSLE disease activities so we chose the objects that could optimally and clearly separate between DA1 and DA3 states and highlight their subgroups. This was done by pruning the misclassified objects from the initial model. The enhanced model showed clearer sub-networks even though the gain in the accuracy was only 10%. While the networks generated here are based on a single gene expression set, multiple lines of evidence from previous SLE studies support their value; whether that be in classifying sub-cluster patient states or indicating possible treatments based on hub genes. It will be important to test the predictive, or replicative, ability of the gene networks to classify additional SLE patient sets, but the permutation analysis conducted here suggests that this should be possible. We believe that machine learning approaches, such as the one demonstrated here, could aid disease understanding and facilitate the clinical and therapeutic stratification of patients. This applies not only to SLE, but to any complex heterogeneous syndrome.

## METHODS

Figure 1, an overview of the analysis pipeline was generated with www.lucidchart.com resources.

### Data and pre-processing

Existing whole blood transcriptome records (Illumina HT-12 V4 bead chip) and clinical metadata from 158 pSLE patients and 48 healthy controls were downloaded (NCBI GEO: GSE65391)^9^ and the values corresponding to DA1, DA3 and control visits extracted. In this analysis, the transcriptome generated per visit to the clinic, and not per patient lifetime, was considered. As such, an individual may be represented in the analysis multiple times (between 1 and 15 times) if their disease status at the time was classified as DA1 or DA3 (Supplementary Fig S12 online). For expression data, gene loci represented by more than one probe were combined and averaged, before each gene locus was log transformed. Batch effects were identified (Variance Partition R package^39^) and corrected (SVA R package). The batch effects identified here were limited to the reported batch replicates from the original metadata (batch 1 and 2) and not found for other phenotypes (Supplementary Fig S13 online).

### Machine learning rule-based modelling to obtain explainable classifiers for DA state

For methodological context, we applied an interpretable learning method based on rough sets that offers classification transparency^11,12^. Given data in the form of a decision table, where rows represent observations and columns are features with the last column being the outcome or decision, rough set algorithms select minimal subsets of features that preserve discernibility between the outcomes for the observations. These subsets of features are called reducts, and are used to generate IF-THEN rules by overlaying them on the observations. An IF-THEN rule consists of the condition part, often called the left-hand side, and the THEN part is the decision given by the rule and often called the right side of the rule. The elements of the IF-part are called descriptors, and are in the form of pairs, feature and its value. To aid interpretation, the rules generated by the model were visualized as RNs, where the nodes are descriptors. For every pair of descriptors in a rule of the RBM, an edge connecting the corresponding nodes is added to the network.

First, expression values were subject to data discretisation, since R. ROSETTA^13^ generates rules for that data form. For each gene, the control data expression mean (μ) and standard deviation (σ) were calculated, and then all DA data for that gene projected onto this threshold frame and discretised (Low ≤ μ − 2σ < Medium > High ≥ μ − 2σ; Numeric values 1, 2, 3).

To generate the initial model, data was first collected into a decision table where unique visit identifiers were the objects and put in rows (n=629), while genes (n=33,006) were variables and constituted columns. The objects were labelled with disease activity, DA1 or DA3, accordingly. Next, Monte Carlo Feature selection (MCFS) algorithm^14^ was applied to obtain a ranked list of informative features with respect to classifying the objects. A significance cut-off for selecting features from the ranked list was obtained by a permutation test (p-value ≤ 0.05). Feature boosting was applied to select the optimal number of features to build the model and then the rule model was visualized with the VisuNet R package^41^.

The initial rule-based model defined above was used as a base to further improve classification. Data (DA1 or DA3 visits) that did not match the left-hand side of any significant rules in the previous model were removed (p-value < 0.05). The MCFS^14^ process was then repeated after object removal. Prior to building the enhanced rule-based model, iterative computational rounds were performed (Feature boosting in Figure 1) in order to select the optimal number of features for building the final predictive model. The significant features from MCFS output were incrementally added to build several rule-based models. The selected features that were used to build the model with the best overall accuracy where chosen for building the final enhanced model using R.ROSETTA^13^ and then visualized using VisuNet^41^.

In order to identify patient subgroups, a matrix was constructed with maintained observations (visits) as rows and rules as columns. The cells for all observations that supported a rule were all assigned 1 or otherwise 0. Hierarchical clustering based on binary distance as the distance function was applied on this matrix.

### Correlating clusters to clinical and phenotypic data

Available metadata, including continuous and categorical clinical values (Supplementary Table S3), were accessed^9^. For continuous variables, a one-way ANOVA following a post-hoc Tukey HSD test was used to compute significance. A Fisher’s exact test was used for the assessment of categorical variables to sub-clusters.

### Correlating rules associated with clusters to clinical and phenotypic data

Empirical values were used to determine the minimal threshold for rule membership to clusters. Rules were considered associated with a cluster if they had a support set matching at least 10% of the cluster’s support set (i.e., observations associated with a cluster; Supplementary Fig S14 online). The association between a cluster’s supported rules and clinical phenotypes was assessed by contrasting phenotype values for supported samples of each rule versus the non-supported samples (categorical variables, non-parametric Wilcoxon test; binary variables, Fisher’s exact test). Supplementary Fig S15 online illustrates this process.

### Model validation

The decision label (DA1 or DA3) was permuted 1,000 times and rule-based models were created for these random sets. A normal distribution was built for the model accuracies and an alpha of 0.05 and a 95% confidence interval used to determine the significance of the p-value. The mean, standard deviation and the standard error for the normal distribution were computed. The accuracy of the original model was compared to the mean μ and standard error σ. If the accuracy of the original model was smaller than μ -σ or greater than μ +σ then the p-value in this case was < 0.05.

### Gene enrichment analysis

Overrepresentation of gene sets belonging to each cluster and the gene sets belonging to rules in DA1 and DA3 were determined using the R package clusterProfiler^42^. The background list was set as initial set of 33,006 available loci.

## ACKNOWLEDGMENTS

We would like to thank Mateusz Garbulowski for providing the rule networks figure under “Disease subtype discovery” in Figure 1 in addition to R. ROSETTA and VisuNet logos.

## CONTRIBUTIONS

J.K. conceptualized the experiment with design input from C.F.B., J.R.S.M., and S.A.Y. J.R.S.M. and S.A.Y. wrote the manuscript. P.S. created the initial model and A.A. enhanced R scripts. K.D. refined the permutation test, implemented the use of binary distance for hierarchal clustering and enhanced the pruning steps for the enhanced model. C.F.B. designed the cluster to phenotype enrichment analysis. S.A.Y. performed all remaining modelling and intersection with phenotypic data. C.F.B., J.R.S.M. and S.A.Y. interpreted the results. J.R.S.M. developed the discussion. L.H. refined manuscript text. S.A.Y. generated all figures and tables with support from J.R.S.M for Figures 3 and 4, and all Supplementary Tables. All authors reviewed the manuscript.

## REFRENCES

1. Kamphuis S, Silverman ED. Prevalence and burden of pediatric-onset systemic lupus erythematosus. Nat Rev Rheumatol. 2010;6(9):538–546. doi:10.1038/nrrheum.2010.121

2. Tsokos GC. Systemic Lupus Erythematosus. N Engl J Med. 2011;365(22):2110–2121. doi:10.1056/NEJMra1100359

3. Peter H. Schur B C. Derivation and validation of the Systemic Lupus International Collaborating Clinics classification criteria for systemic lupus erythematosus - Petri - 2012 - Arthritis & Rheumatism - Wiley Online Library. Accessed March 2, 2021. https://onlinelibrary.wiley.com/doi/full/10.1002/art.34473

4. Bombardier C, Gladman DD, Urowitz MB, et al. Derivation of the sledai. A disease activity index for lupus patients. Arthritis Rheum. 1992;35(6):630–640. doi:10.1002/art.1780350606

5. Li Q-Z, Zhou J, Lian Y, et al. Interferon signature gene expression is correlated with autoantibody profiles in patients with incomplete lupus syndromes. Clin Exp Immunol. 2010;159(3):281–291. doi:10.1111/j.1365-2249.2009.04057.x

6. Kyogoku C, Smiljanovic B, Grün JR, et al. Cell-Specific Type I IFN Signatures in Autoimmunity and Viral Infection: What Makes the Difference? PLOS ONE. 2014;8(12):e83776. doi:10.1371/journal.pone.0083776

7. Demirkaya E, Sahin S, Romano M, Zhou Q, Aksentijevich I. New Horizons in the Genetic Etiology of Systemic Lupus Erythematosus and Lupus-Like Disease: Monogenic Lupus and Beyond. J Clin Med. 2020;9(3):712. doi:10.3390/jcm9030712

8. Marion TN, Postlethwaite AE. Chance, genetics, and the heterogeneity of disease and pathogenesis in systemic lupus erythematosus. Semin Immunopathol. 2014;36(5):495–517. doi:10.1007/s00281-014-0440-x

9. Banchereau R, Hong S, Cantarel B, et al. Personalized Immunomonitoring Uncovers Molecular Networks that Stratify Lupus Patients. Cell. 2016;165(3):551–565. doi:10.1016/j.cell.2016.03.008

10. Mirza B, Wang W, Wang J, Choi H, Chung NC, Ping P. Machine Learning and Integrative Analysis of Biomedical Big Data. Genes. 2019;10(2):87. doi:10.3390/genes10020087

11. Komorowski J. 6.02 - Learning Rule-Based Models - The Rough Set Approach. In: Brahme A, ed. Comprehensive Biomedical Physics. Elsevier; 2014:19–39. doi:10.1016/B978-0-444-53632-7.01102-3

12. Skowron A, Dutta S. Rough sets: past, present, and future. Nat Comput. 2018;17(4):855–876. doi:10.1007/s11047-018-9700-3

13. Garbulowski M, Diamanti K, Smolińska K, et al. R.ROSETTA: an interpretable machine learning framework. BMC Bioinformatics. 2021;22(1):110. doi:10.1186/s12859-021-04049-z

14. Dramiński M, Rada-Iglesias A, Enroth S, Wadelius C, Koronacki J, Komorowski J. Monte Carlo feature selection for supervised classification. Bioinformatics. 2008;24(1):110–117. doi:10.1093/bioinformatics/btm486

15. Gao G, Zhu C, Liu E, Nabi IR. Reticulon and CLIMP-63 regulate nanodomain organization of peripheral ER tubules. PLoS Biol. 2019;17(8):e3000355–e3000355. doi:10.1371/journal.pbio.3000355

16. Xiahou Z, Wang X, Shen J, et al. NMI and IFP35 serve as proinflammatory DAMPs during cellular infection and injury. Nat Commun. 2017;8. doi:10.1038/s41467-017-00930-9

17. Bradley SJ, Suarez-Fueyo A, Moss DR, Kyttaris VC, Tsokos GC. T Cell Transcriptomes Describe Patient Subtypes in Systemic Lupus Erythematosus. PloS One. 2015;10(11):e0141171. doi:10.1371/journal.pone.0141171

18. Harley JB, Alarcón-Riquelme ME, Criswell LA, et al. Genome-wide association scan in women with systemic lupus erythematosus identifies susceptibility variants in ITGAM, PXK, KIAA1542 and other loci. Nat Genet. 2008;40(2):204–210. doi:10.1038/ng.81

19. Löfgren SE, Frostegård J, Truedsson L, et al. Genetic association of miRNA-146a with systemic lupus erythematosus in Europeans through decreased expression of the gene. Genes Immun. 2012;13(3):268–274. doi:10.1038/gene.2011.84

20. Toro-Domínguez D, Martorell-Marugán J, Goldman D, Petri M, Carmona-Sáez P, Alarcón-Riquelme ME. Stratification of Systemic Lupus Erythematosus Patients Into Three Groups of Disease Activity Progression According to Longitudinal Gene Expression. Arthritis Rheumatol. 2018;70(12):2025–2035. doi:https://doi.org/10.1002/art.40653

21. Tesser A, de Carvalho LM, Sandrin-Garcia P, et al. Higher interferon score and normal complement levels may identify a distinct clinical subset in children with systemic lupus erythematosus. Arthritis Res Ther. 2020;22(1):91. doi:10.1186/s13075-020-02161-8

22. Han BK, Wysham KD, Cain KC, Tyden H, Bengtsson AA, Lood C. Neutrophil and lymphocyte counts are associated with different immunopathological mechanisms in systemic lupus erythematosus. Lupus Sci Med. 2020;7(1). doi:10.1136/lupus-2020-000382

23. Greene CS, Tan J, Ung M, Moore JH, Cheng C. Big Data Bioinformatics. J Cell Physiol. 2014;229(12):1896–1900. doi:10.1002/jcp.24662

24. Azodi CB, Tang J, Shiu S-H. Opening the Black Box: Interpretable Machine Learning for Geneticists. Trends Genet. 2020;36(6):442–455. doi:10.1016/j.tig.2020.03.005

25. Glaab E, Bacardit J, Garibaldi JM, Krasnogor N. Using Rule-Based Machine Learning for Candidate Disease Gene Prioritization and Sample Classification of Cancer Gene Expression Data. PLOS ONE. 2012;7(7):e39932. doi:10.1371/journal.pone.0039932

26. Bange FC, Vogel U, Flohr T, Kiekenbeck M, Denecke B, Böttger EC. IFP 35 is an interferon-induced leucine zipper protein that undergoes interferon-regulated cellular redistribution. J Biol Chem. 1994;269(2):1091–1098.

27. Das A, Dinh PX, Panda D, Pattnaik AK. Interferon-Inducible Protein IFI35 Negatively Regulates RIG-I Antiviral Signaling and Supports Vesicular Stomatitis Virus Replication. J Virol. 2014;88(6):3103–3113. doi:10.1128/JVI.03202-13

28. Lin Y-L, Lin S-C. Analysis of the CD161-expressing cell quantities and CD161 expression levels in peripheral blood natural killer and T cells of systemic lupus erythematosus patients. Clin Exp Med. 2017;17(1):101–109. doi:10.1007/s10238-015-0402-1

29. Kurioka A, Cosgrove C, Simoni Y, et al. CD161 Defines a Functionally Distinct Subset of Pro-Inflammatory Natural Killer Cells. Front Immunol. 2018;9. doi:10.3389/fimmu.2018.00486

30. Gao L, Wang Q, Ren W, et al. The RBP1-CKAP4 axis activates oncogenic autophagy and promotes cancer progression in oral squamous cell carcinoma. Cell Death Dis. 2020;11(6):488. doi:10.1038/s41419-020-2693-8

31. Clarke AJ, Ellinghaus U, Cortini A, et al. Autophagy is activated in systemic lupus erythematosus and required for plasmablast development. Ann Rheum Dis. 2015;74(5):912–920. doi:10.1136/annrheumdis-2013-204343

32. Zhong Z, Umemura A, Sanchez-Lopez E, et al. NF-κB Restricts Inflammasome Activation via Elimination of Damaged Mitochondria. Cell. 2016;164(5):896–910. doi:10.1016/j.cell.2015.12.057

33. Kikuchi A, Fumoto K, Kimura H. The Dickkopf1-cytoskeleton-associated protein 4 axis creates a novel signalling pathway and may represent a molecular target for cancer therapy. Br J Pharmacol. 2017;174(24):4651–4665. doi:10.1111/bph.13863

34. Mackay M, Oswald M, Sanchez-Guerrero J, et al. Molecular signatures in systemic lupus erythematosus: distinction between disease flare and infection. Lupus Sci Amp Med. 2016;3(1):e000159. doi:10.1136/lupus-2016-000159

35. Yalcin E, Beker MC, Turkseven S, et al. Evidence that melatonin downregulates Nedd4-1 E3 ligase and its role in cellular survival. Toxicol Appl Pharmacol. 2019;379:114686. doi:10.1016/j.taap.2019.114686

36. Bruck R, Aeed H, Avni Y, et al. Melatonin inhibits nuclear factor kappa B activation and oxidative stress and protects against thioacetamide induced liver damage in rats. J Hepatol. 2004;40(1):86–93. doi:10.1016/S0168-8278(03)00504-X

37. Huang H, Liu X, Chen D, et al. Corrigendum to “Melatonin prevents endothelial dysfunction in SLE by activating the nuclear receptor retinoic acid-related orphan receptor-α” [Int. Immunopharmacol. 83 (2020) 106365]. Int Immunopharmacol. 2020;86:106817. doi:10.1016/j.intimp.2020.106817

38. Bonomini F, Dos Santos M, Veronese FV, Rezzani R. NLRP3 Inflammasome Modulation by Melatonin Supplementation in Chronic Pristane-Induced Lupus Nephritis. Int J Mol Sci. 2019;20(14):3466. doi:10.3390/ijms20143466

39. Hoffman GE, Schadt EE. variancePartition: interpreting drivers of variation in complex gene expression studies. BMC Bioinformatics. 2016;17(1):483. doi:10.1186/s12859-016-1323-z

40. Leek JT, Johnson WE, Parker HS, Jaffe AE, Storey JD. The sva package for removing batch effects and other unwanted variation in high-throughput experiments. Bioinforma Oxf Engl. 2012;28(6):882–883. doi:10.1093/bioinformatics/bts034

41. Lab KB. Komorowskilab/VisuNet.; 2020. Accessed March 2, 2021. https://github.com/komorowskilab/VisuNet

42. Yu G, Wang L-G, Han Y, He Q-Y. clusterProfiler: an R Package for Comparing Biological Themes Among Gene Clusters. OMICS J Integr Biol. 2012;16(5):284–287. doi:10.1089/omi.2011.0118

