## Supplementary figures for "Interpretable machine learning identifies paediatric Systemic Lupus Erythematosus subtypes based on gene expression data"

**ORCID**: J.R.S.M., 0000-0002-0850-230X; S.A.Y.,0000-0002-7201-2604; L.H., 0000-0003-4140-3423; K.D., 0000-0002-4922-8415; J.K., 0000-0002-0766-8789

[+]Equal contribution
[*]Corresponding authors

### SUPPLEMENTARY FIGURES

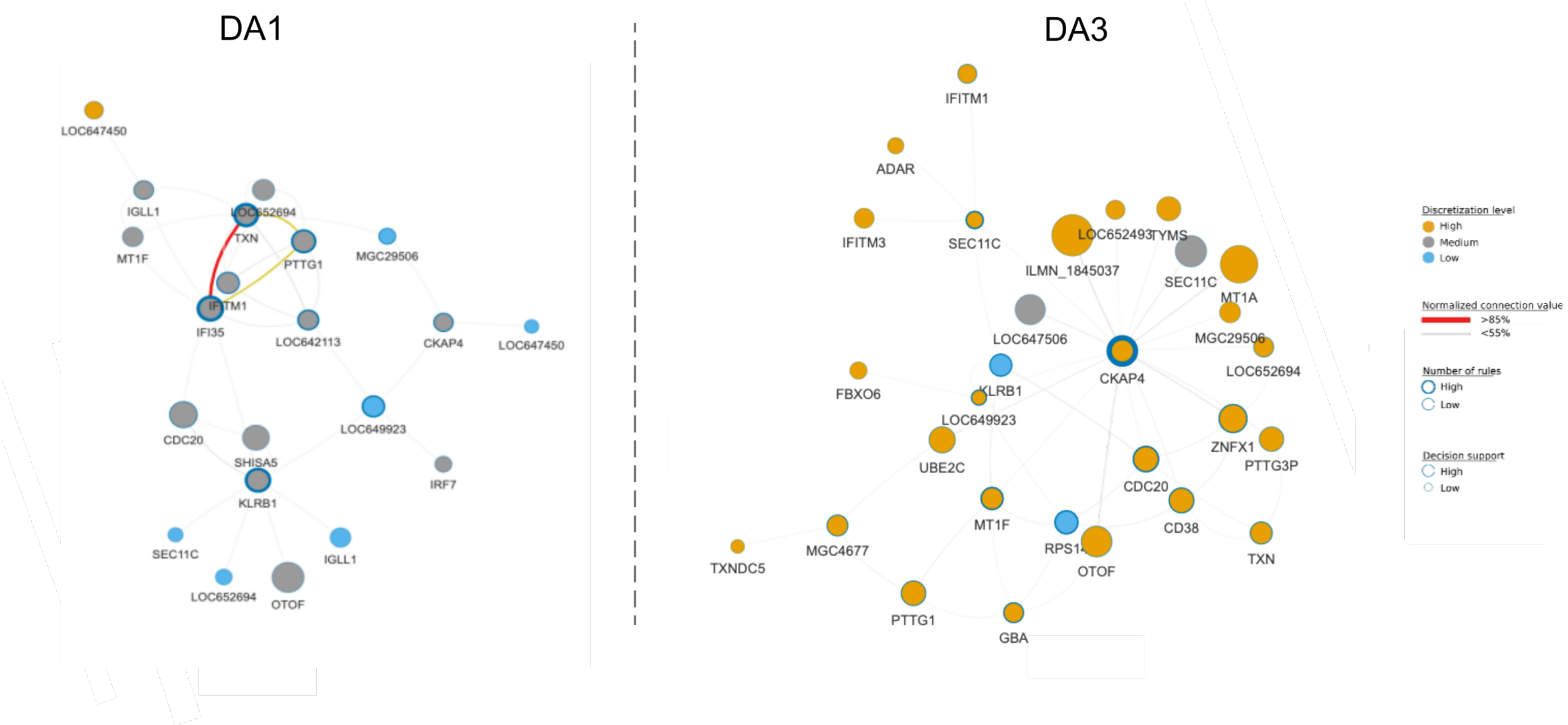

**Supplementary Figure S1.**
The initial rule-based model visualised as a rule network distinguishes pSLE disease states DA1 and DA3. Discretised gene expression value is indicated by the colour of the node circles (high, medium, low; orange, grey, blue). Node size is proportional to the number of objects that support rules for a decision class (node circle size), node border is proportional to the number of rules associated to a node (low, high; circle border thin, thick) and lines connecting nodes are normalised connection values (<55%, ≥85%; grey, red with increasing line thickness per support interval). The latter represent the strength of co-appearance for the connected nodes in rules supporting a decision class.

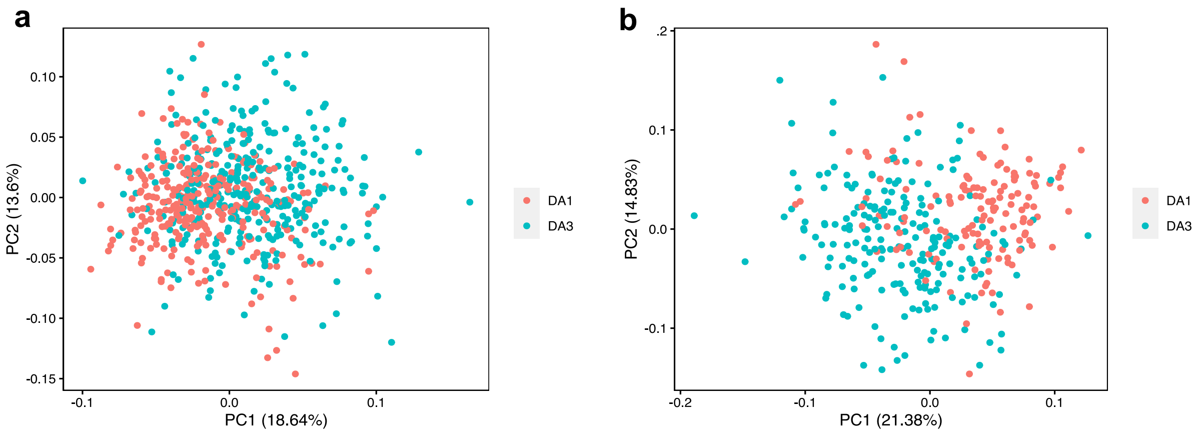

**Supplementary Figure S2.**
Principal component analysis (PCA) of the initial rule-based model **(a)** before and **(b)** after pruning misclassified DA1 or DA3 observations.

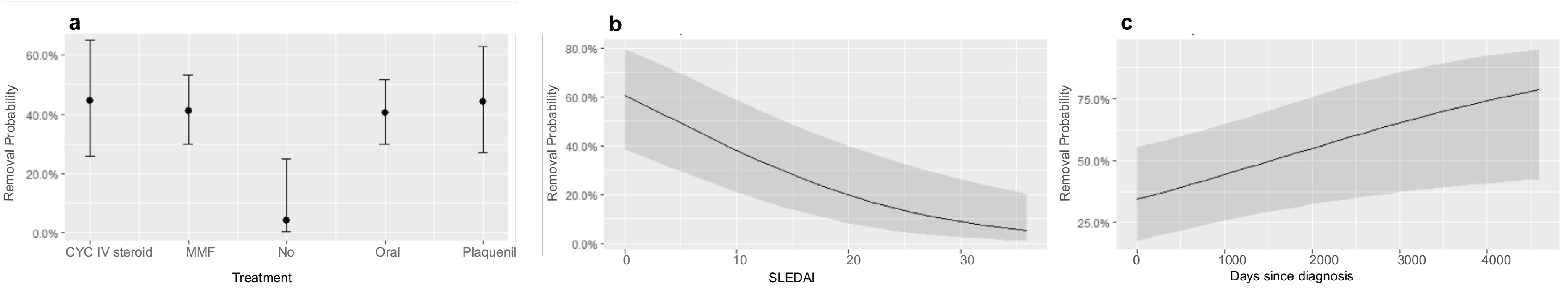

**Supplementary Figure S3.**
Relationship between the probability of pruning observations and phenotypic measure for **(a)** treatment prescribed to patients, **(b)** SLEDAI score and **(c)** number of days since diagnosis

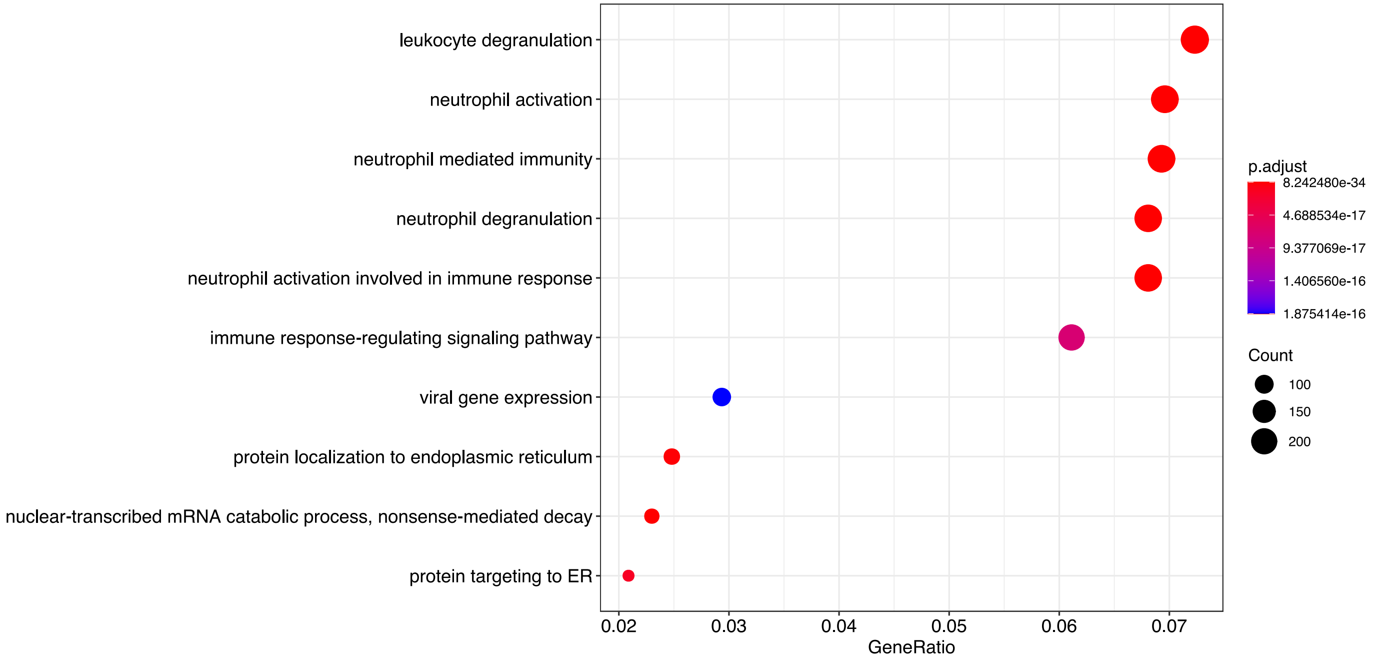

**Supplementary Figure S4.**
Enrichment of gene ontology biological process terms based on 4,980 genes from the pruned DA1 and DA3 dataset.

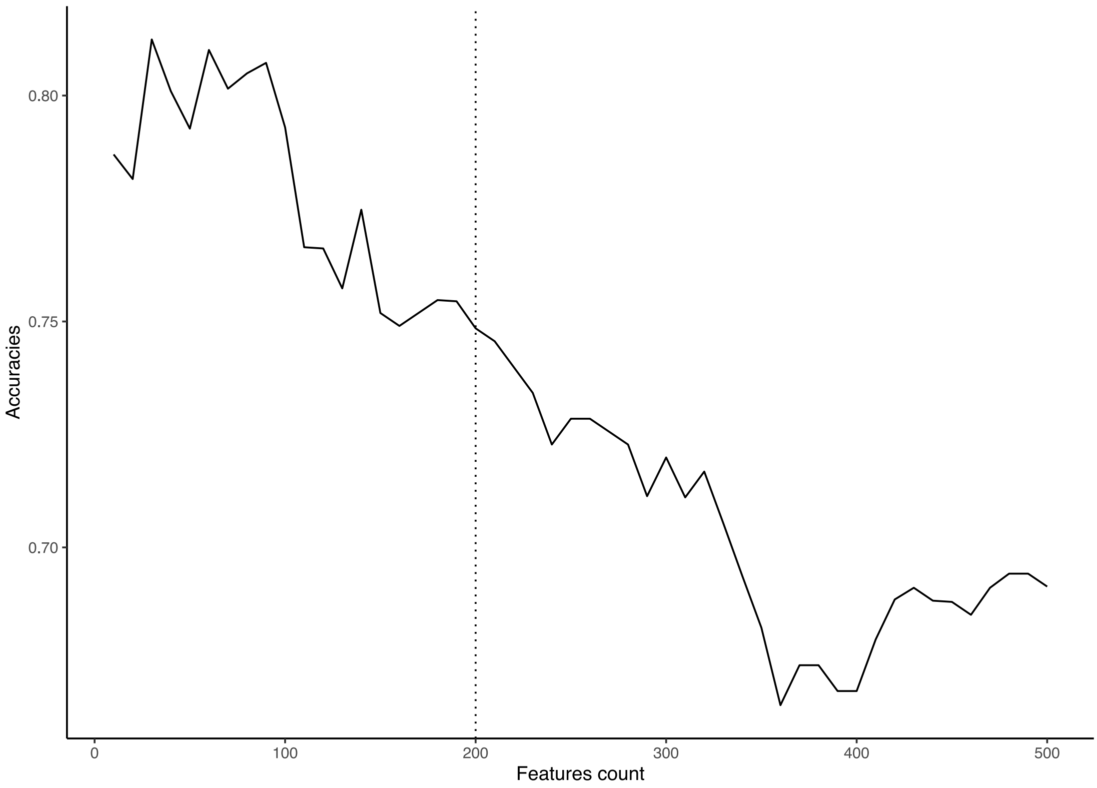

**Supplementary Figure S5.**
Feature boosting shows that the accuracy of the model drops after using the first 200 features to build the rule-based model

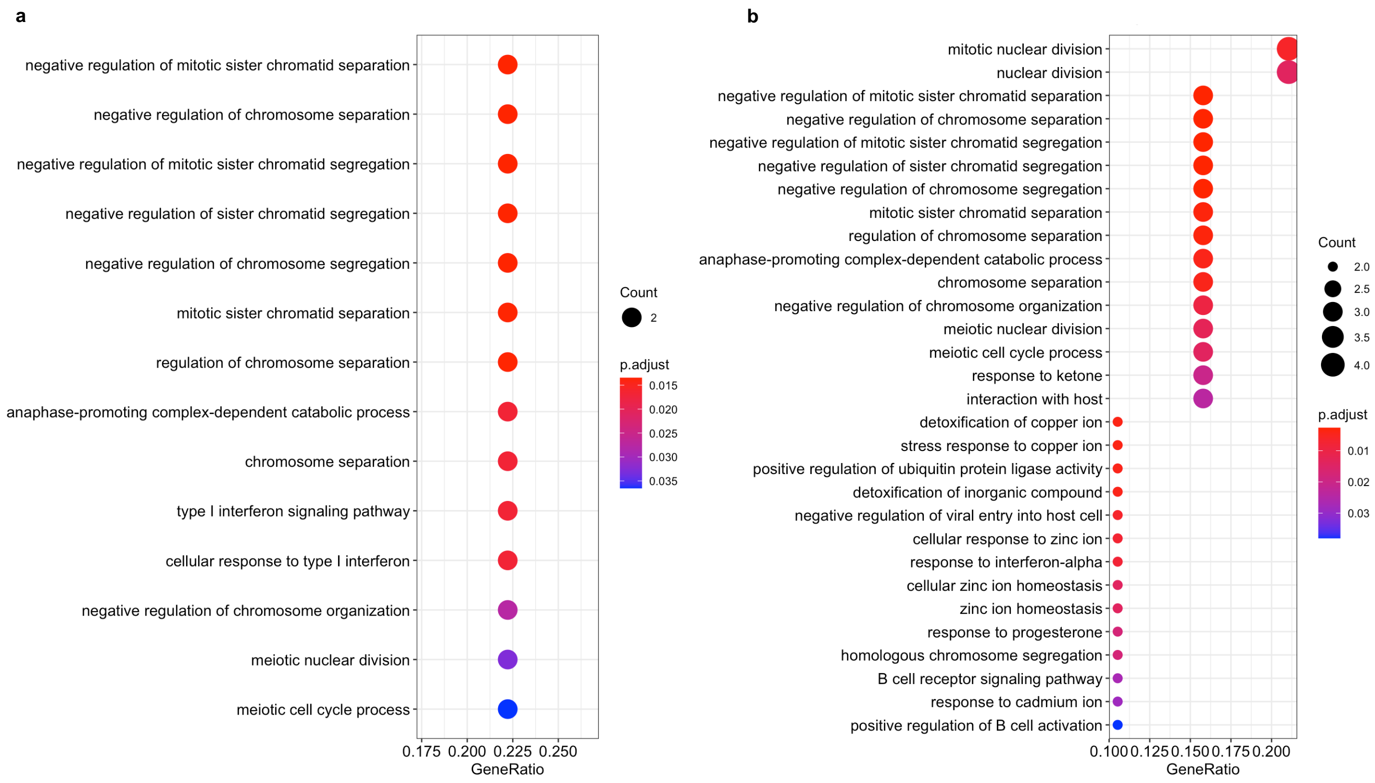

**Supplementary Figure S6.**
Graphical enrichment of Gene Ontology biological process terms based on the enhanced model gene sets for **(a)** DA1 and **(b)** DA3 (12 and 21 loci respectively).

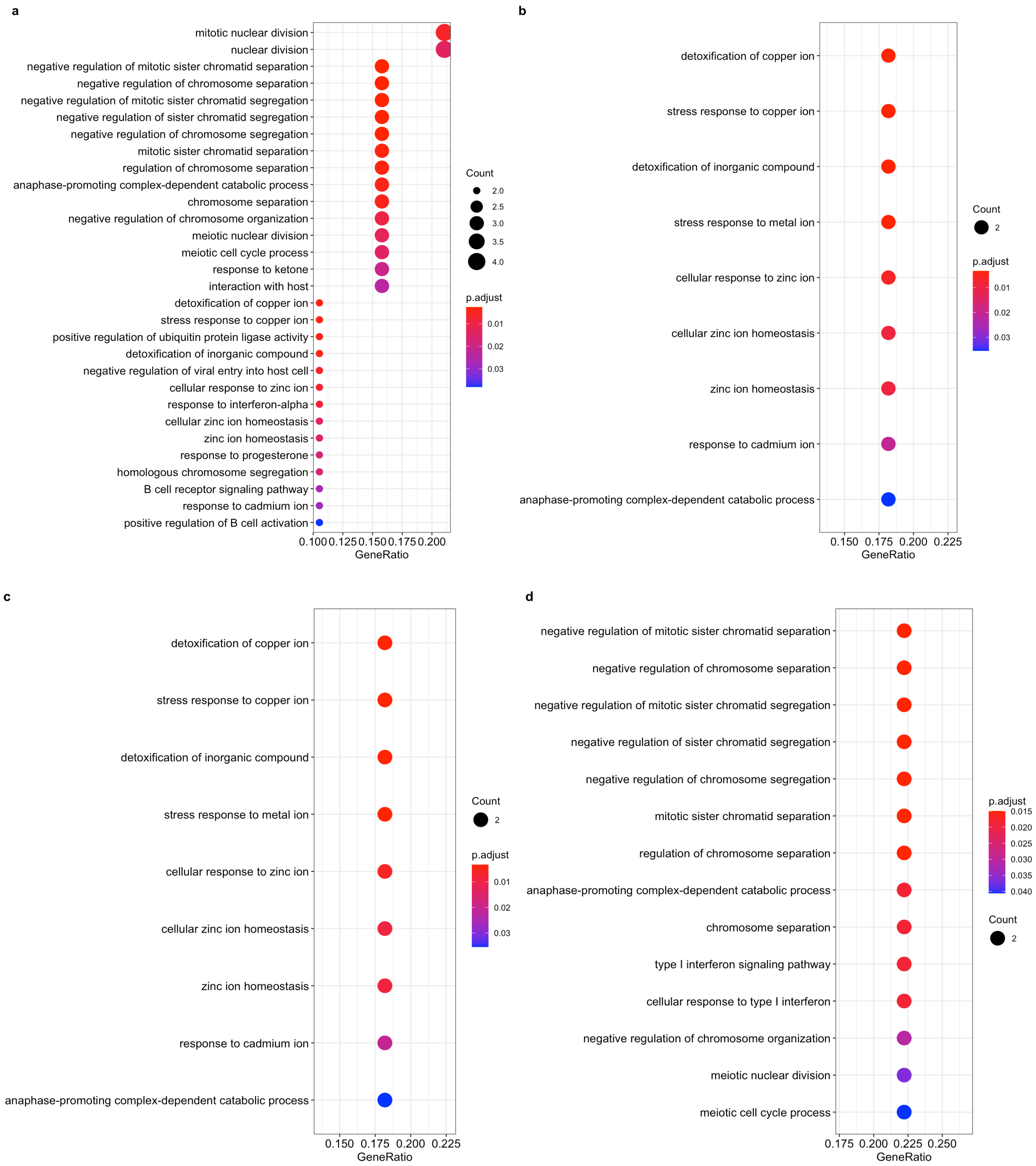

**Supplementary Figure S7.**
Graphical enrichment of Gene Ontology biological process terms based on the five sub-clusters **(a)** C3, **(b)** C5, **(c)** C4 and **(d)** C1 determined via hierarchical analyses of DA1 and DA. C2 did not have any enriched terms.

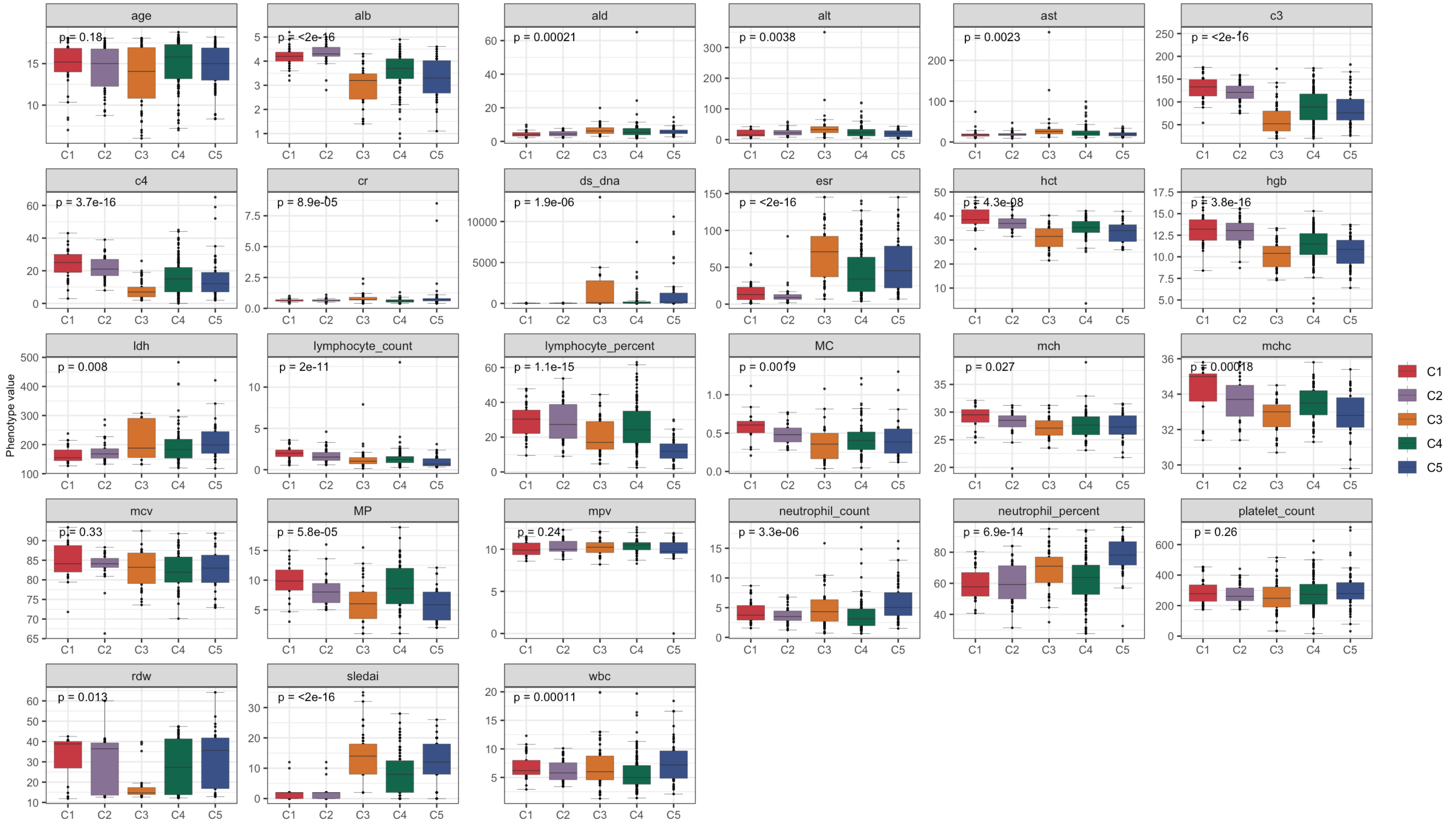

**Supplementary Figure S8.**
Boxplots illustrating the distribution of all 27 continuous clinical values correlated with clusters. Phenotype abbreviations are shown in Supplementary Table S3.

a

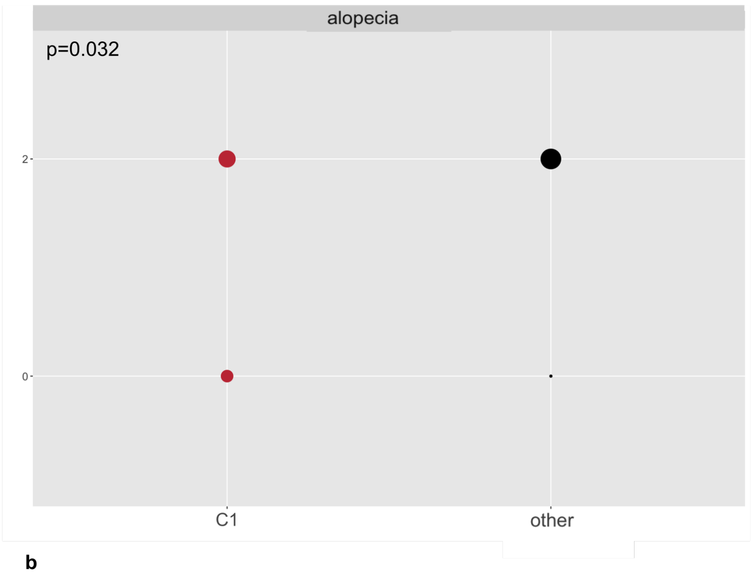

b

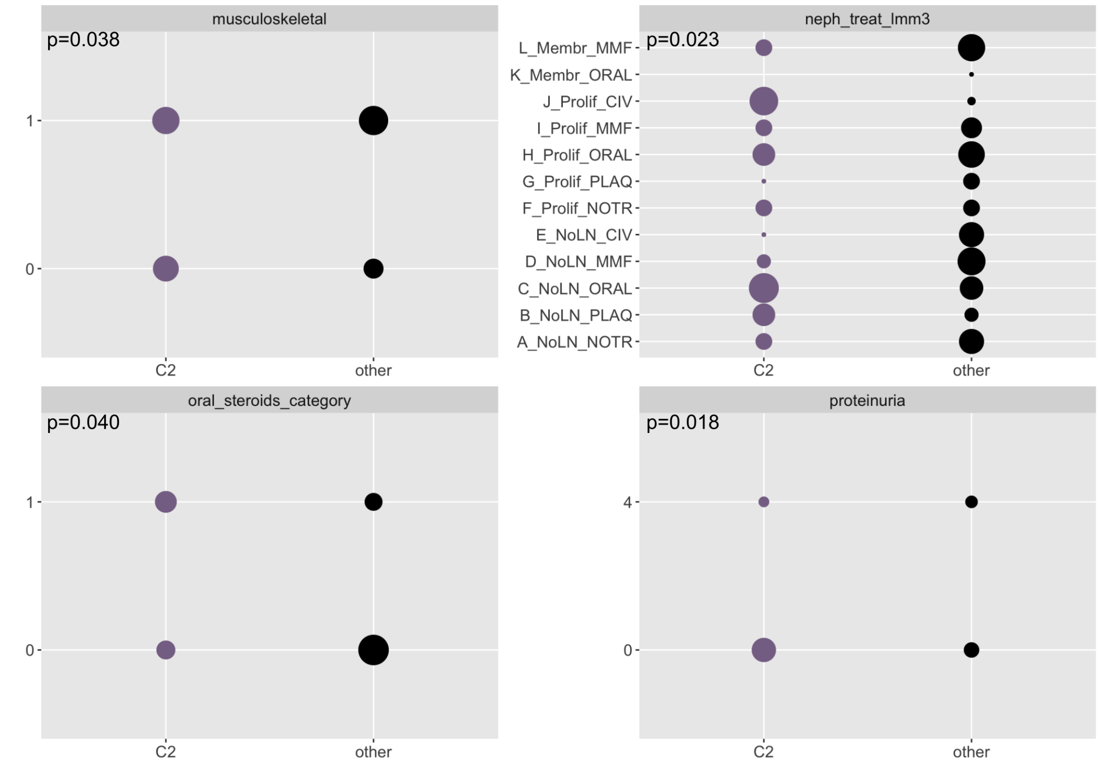

**Supplementary Figure S9.**
Balloon plots illustrating the distribution of the significant variables for **(a)** C1 and **(b)** C2 (p-value ≤ 0.05).

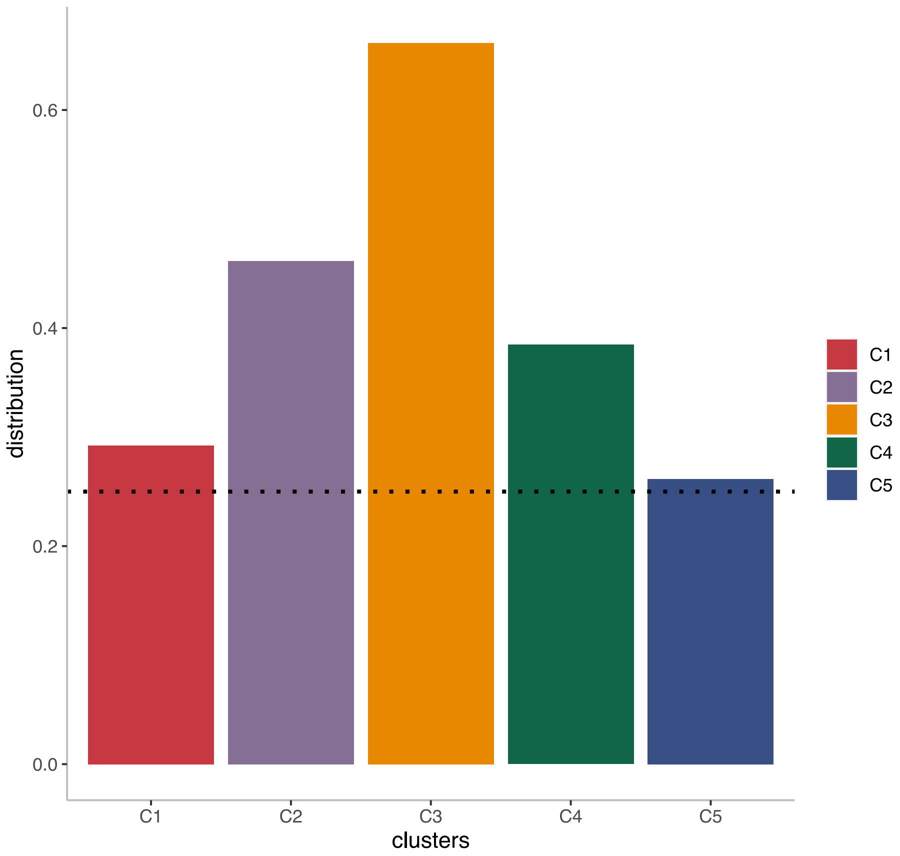

**Supplementary Figure S10.**
Frequency distribution for all rules with support set matching at least 10% of the patient visits assigned to each of the discovered clusters. A threshold of 20% was chosen to associate rules to clusters.

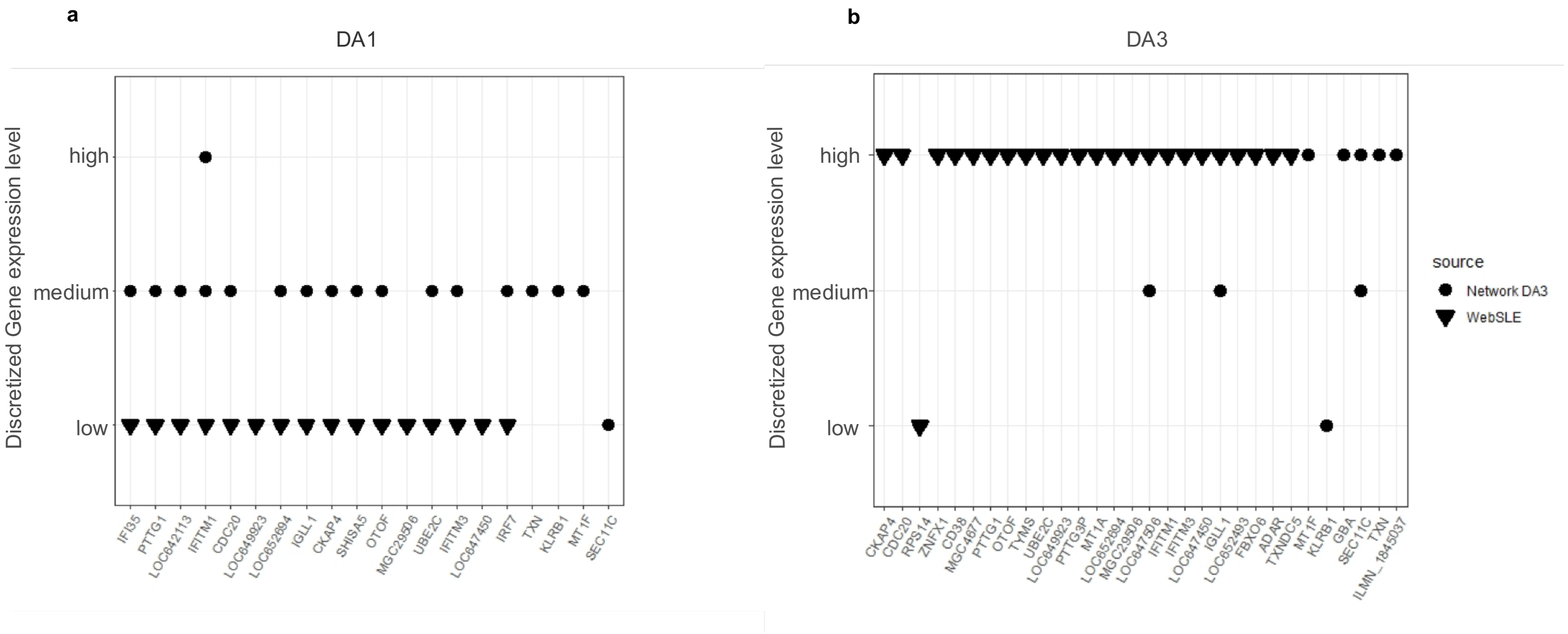

**Supplementary Figure S11.**

Comparison between the genes reported by the enhanced rule model and those that appeared in the Differential Gene Expression (DGE) analysis from the original WebSLE analysis of Banchereau *et al*., 2016 for **(a)** DA1 and **(b)** DA3. The x axis shows the genes discovered by the rules and the y axis indicated the state of the gene. Genes such as *TXN* were key to the rule based model, but not noted in the DGE analysis.

Banchereau R, Hong S, Cantarel B, et al. Personalized Immunomonitoring Uncovers Molecular Networks that Stratify Lupus Patients. *Cell*. 2016;165(3):551-565. doi:10.1016/j.cell.2016.03.008

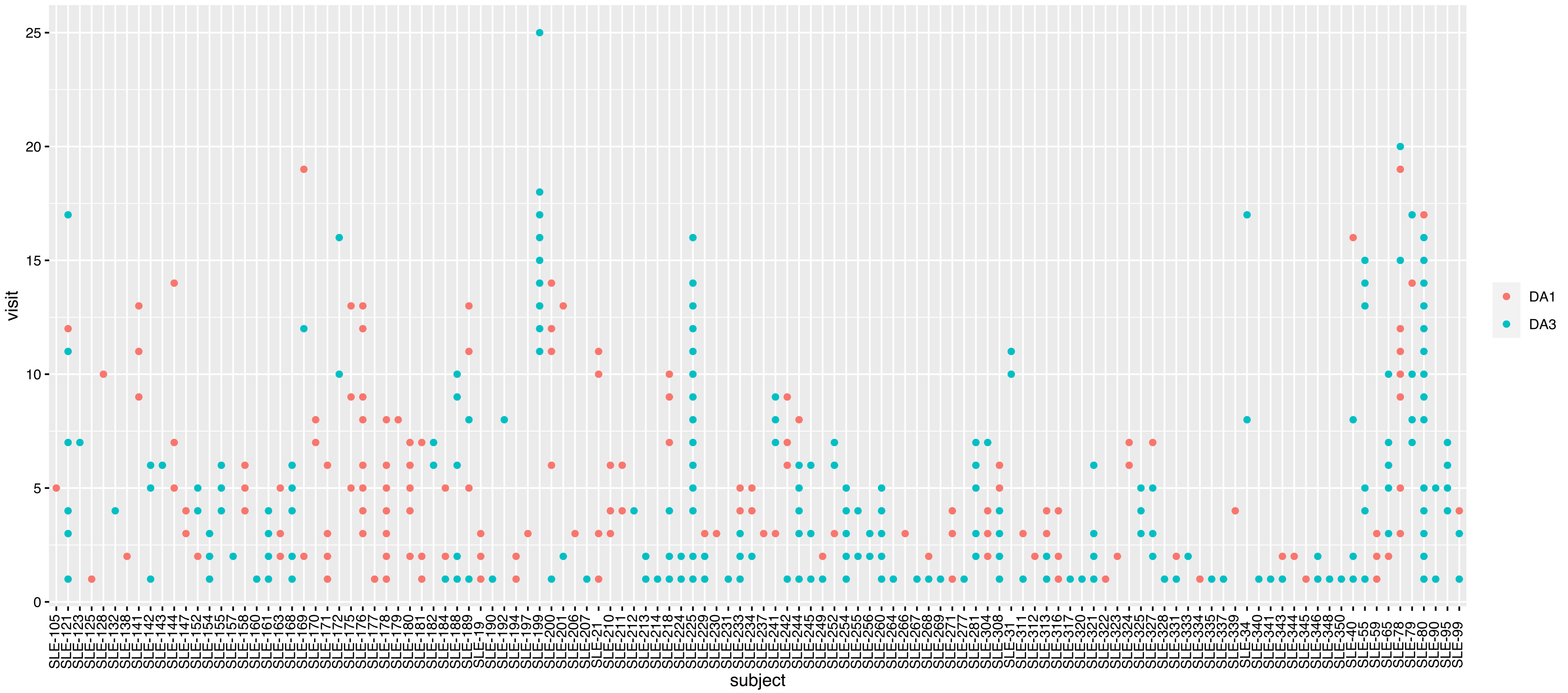

**Supplementary Figure S12.**
The number of visits per patient and their DA class at that time were recorded. Visits with missing data or a DA not equal to 1 or 3 were removed from the original dataset of Banchereau *et al*., 2016.
Banchereau R, Hong S, Cantarel B, et al. Personalized Immunomonitoring Uncovers Molecular Networks that Stratify Lupus Patients. *Cell*. 2016;165(3):551-565. doi:10.1016/j.cell.2016.03.008

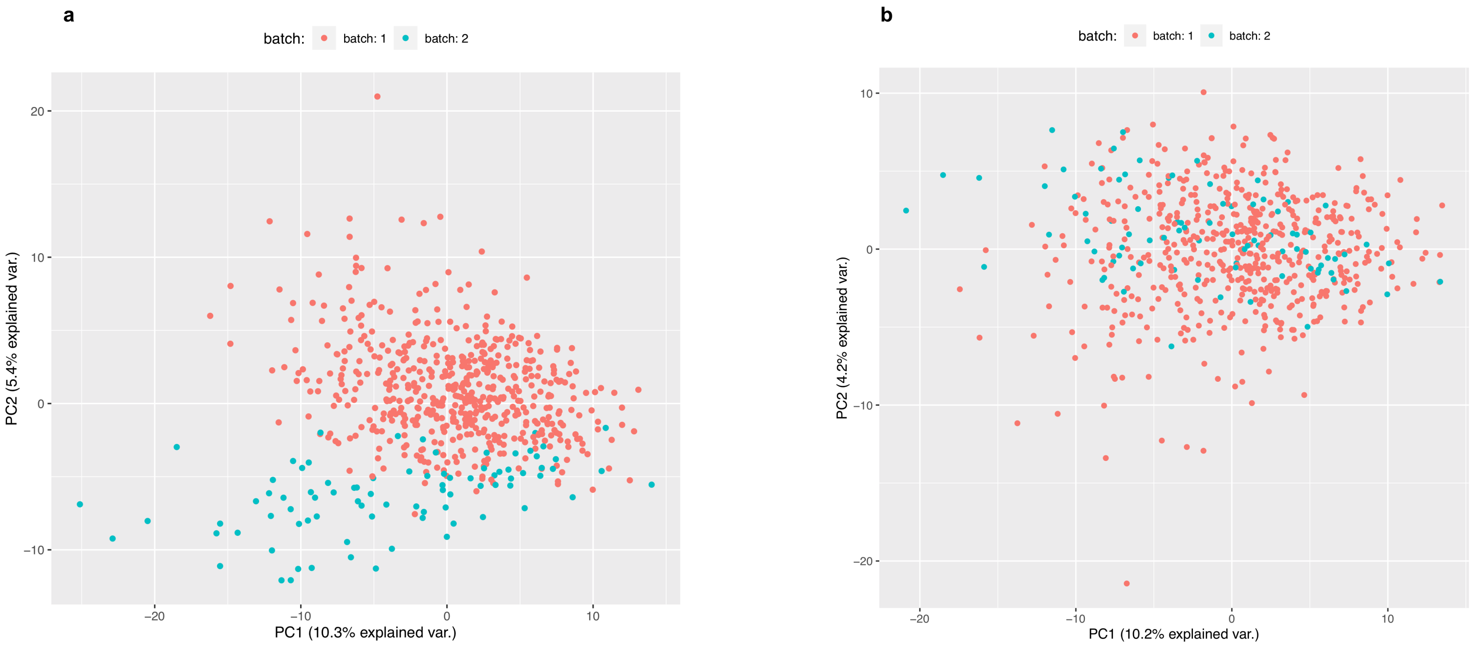

**Supplementary Figure S13.**
Principal component analysis plot for the DA1 and DA3 visits **(a)** before and **(b)** after batch effect correction.

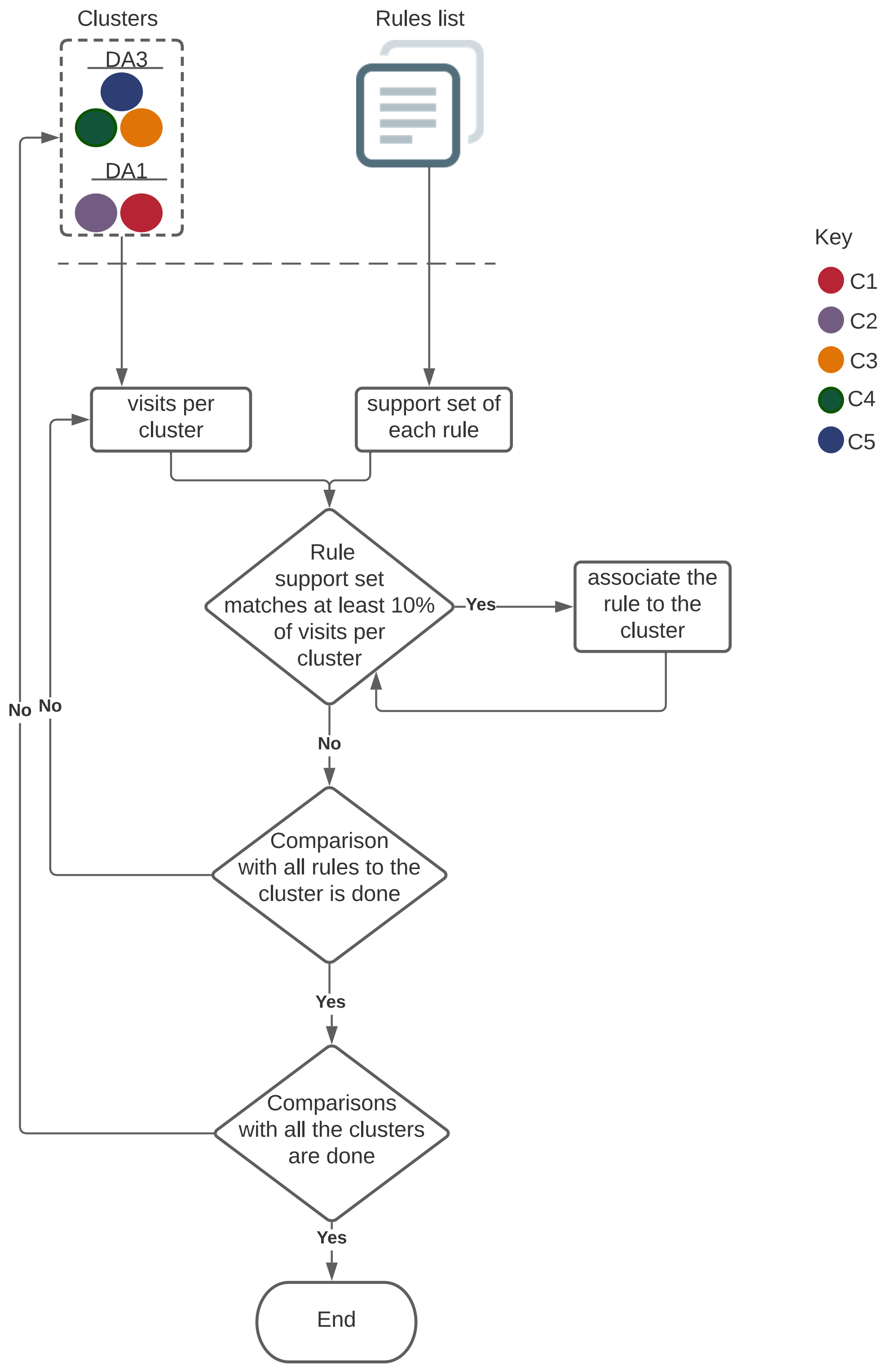

**Supplementary Figure S14.**
Flow chart illustrating the process of associating rules with each of the five discovered clusters.

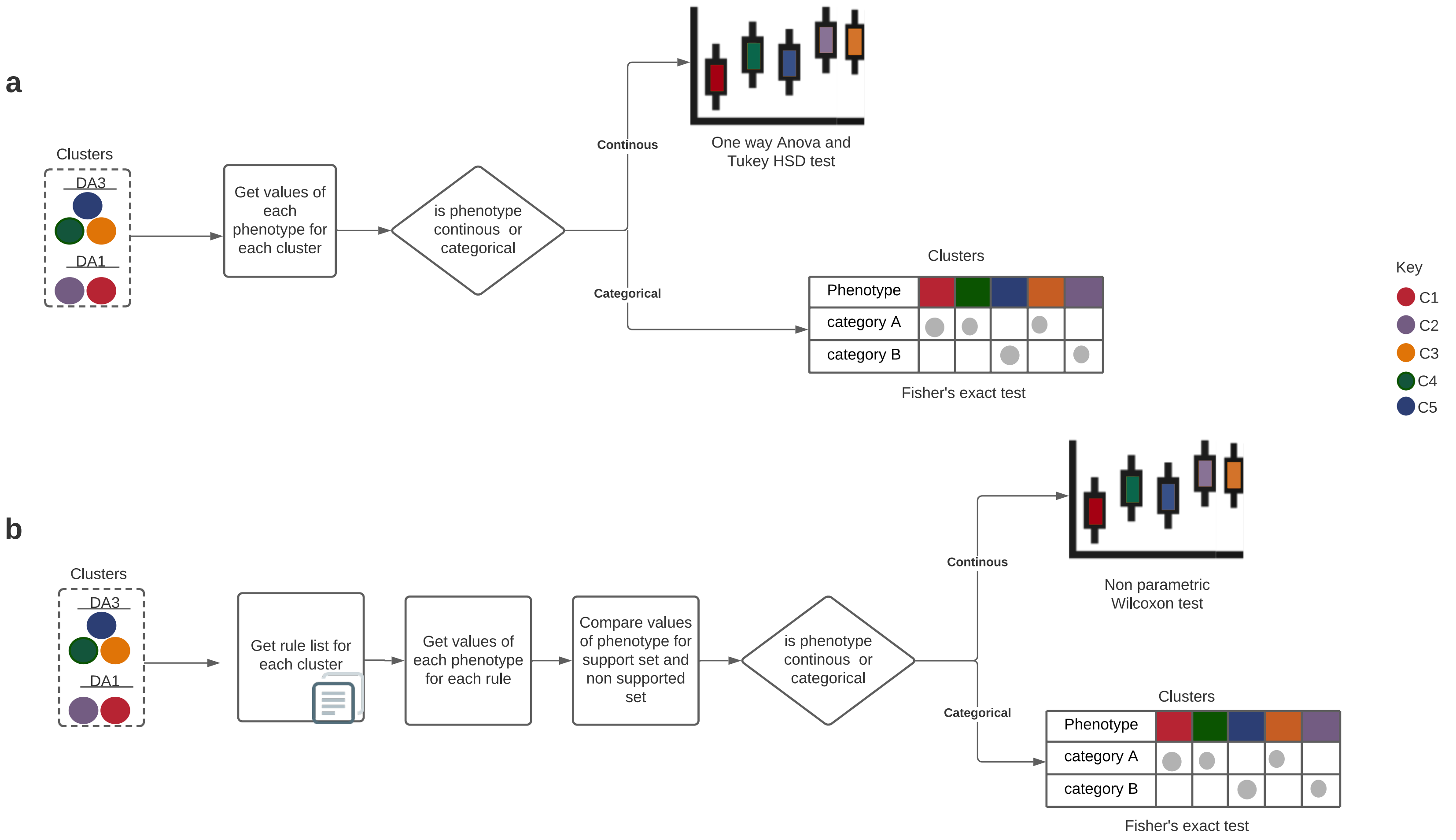

**Supplementary Figure S15.**
Flow chart for significance calculation of **(a)** continuous phenotypes/clinical variables, and **(b)** categorical phenotypes/clinical variables per each rule associated with the five discovered clusters.

**SUPPLEMENTARY TABLES**
All tables are included in excel format

**Supplementary Table S1. Rules, genes and discretised expression value.**

**Supplementary Table S2. Summary of clinical variables with significant difference between at least one cluster pair.**

**Supplementary Table S3. Clinical phenotype abbreviations**

**Supplementary Table S4. Rule-associated continuous clinical phenotypes for each sub-cluster.**

**Supplementary Table S5. Rule-associated categorical clinical phenotypes for each sub-cluster.**
